# Rescuable effect of R-salbutamol in LPS-induced immune dysfunction of sepsis

**DOI:** 10.1101/2021.05.17.444573

**Authors:** Huimin Beng, Shanping Wang, Junhua Hu, Xinglong Liang, Haolong Qin, Wen Tan

## Abstract

Sepsis is a severe life-threatening condition caused by a dysregulated host response to infection. So far, there are no pharmacotherapies to stop sepsis. Salbutamol, a commonly used β_2_-adrenoreceptor agonist, has found to be potential in regulating immune response dysfunction and exert anti-inflammatory effect. However, salbutamol exists two isomers. R-isomer exhibits the therapeutic effect and clinical benefit, while S-isomer proves to be detrimental rather than benign. So, in this study, we investigated the preventive and therapeutic effect of R-salbutamol (R-sal), S-salbutamol (S-sal) or racemic mixture in a mouse model of lipopolysaccharide (LPS)-induced sepsis. Dexamethasone (Dex) was set as comparison. The results showed that R-sal markedly improved seven-day survival rate of septic mice both administered before or after LPS. Whereas Dex showed toxic and accelerated the death of septic mice when given before LPS injection. Lung histological examination and lung function assay revealed that LPS challenge resulted in acute lung damage, including inflammatory cell infiltration, thickened alveolar septa and congestion, and decreased minute volume in septic mice. R-sal pretreatment efficiently inhibited these changes, accompanying by markedly reduced lung MPO level, serum cytokines levels and lactate release and significantly restored the lymphocytes and suppressed the percentage of monocytes. Racemic mixture exhibited diminished effects while S-sal showed enhanced cytokines release. In addition, R-sal pretreatment showed a better improvement in prognostic pulmonary function at day4 in survived mice than that of Rac-sal. Collectively, our results indicate the potential benefit of R-sal for sepsis and sepsis-induced lung injury.

## 1. Introduction

Sepsis is a lift-threatening clinical condition caused by a dysfunction immune response to infection, and is characterized by an overacting systemic inflammation response that followed by anergy, lymphopenia, elevated so-called “cytokine storm” and finally multiorgan dysfunction[1][2]. The deaths due to sepsis remain relatively high worldwide[3]. The most vulnerable organ in the context of sepsis is the lung. Sepsis induces acute lung injury (ALI) and acute respiratory distress syndrome (ARDS), then leads to death[4][5]. As of now, there is no specific pharmacotherapies have been identified. In clinic, the strategies for the treatment of sepsis are removing the infection tissue, initial fluid resuscitation and broad-spectrum antibiotic therapy. However, the indiscriminate of antibiotic can increase antimicrobial resistance. Moreover, even if the patients survived, they are more prone to develop sepsis-induced long-term organ damage and disability[6].

Septic infection could be triggered by ranges of insults, including bacteria, viruses, or fungi. For example, since the end of 2019, the outbreak of coronavirus disease 2019 (COVID-19) pandemic was occurred by severe acute respiratory syndrome coronavirus 2 (SARS-CoV-2) infection. To date, exceed millions of cases worldwide have been reported and the case fatality rate still shows a tendency of climbing high. Hospitalized severe or critically ill patients infected with SARS-CoV-2 to some extent are associated with developing lethal sepsis, which is the leading cause of mortality[7][8][9]. Many therapeutic agents that target antiviral and anti-inflammatory effect are still under active investigation. However, so far, no therapeutic agents have been shown to reduce mortality besides dexamethasone. In the RECOVERY trials, the effectiveness of dexamethasone in lower 28-day mortality is only among those ventilated patients[10]. Moreover, the use of systemic steroids in sepsis is still controversial due to the contradictory results from clinical trials.[11][12][13][14]. Thus, it’s clear that more researches are urgently needed to identify more effective drugs for treating sepsis and sepsis-induced immune dysfunction.

β-adrenergic function is a well-known powerful modulator of the immune responses[15]. β_2_-adrenoreceptor agonists (β_2_-agonists) are the most commonly used bronchodilators to relieve asthma symptoms. Because of the possibility that β_2_-agonists might promote the clearance of alveolar fluid and decrease pulmonary inflammation, the effectiveness of β_2_-agonists has been tested to treat ARDS[16][17][18]. However, in the setting of ARDS, the results of these tests are controversial[19]. Previous studies suggested that, in pre-clinical trials, β_2_-agonists showed success in improving outcomes[20][21]. By contrast, larger trials were terminated on the ground of futility and concerns about safety. In this regard, we found that all the tested β_2_-agonists in the previous clinical trials were in racemic mixture of both R- and S-isomers. However, it’s well known that only R-isomers of β_2_-agonists exhibit the therapeutic effect and clinical benefit, while S-isomers of β_2_-agonists prove to be detrimental rather than benign[22][23]. Hence, we can hypothesize that R-isomers of β_2_-agonists may exhibit a superiority over racemic mixture in terms of sepsis and sepsis-induced lung damage.

Herein, in this study, lipopolysaccharide (LPS), a widely used exogenous toxin, was chosen to induce systemic inflammation in mice to mimic the clinical features of sepsis. R-salbutamol (R-sal), S-salbutamol (S-sal) and the mixture of (R+S)-salbutamol (Rac-sal, the most commonly used β_2_-agonist) were employed to pretreated or treated the mice separately before or after LPS stimulation. As such, dexamethasone was set as comparison. Then we studied their efficacy in improving survival and alleviating systemic inflammation in terms of this mouse model of experimental sepsis, mainly observing: i) acute survival rate; ii) lung histopathological damage; iii) MPO levels; iv) hematology examinations; v) inflammatory cytokines; vi) prognostic lung function.

## 2. Materials and methods

### Reagents

R-salbutamol sulfate (R-sal, >99% purity, 99.85% ee) and S-salbutamol sulfate (S-sal, >99% purity, 92.73% ee) were obtained from Suzhou Junning R&D Center for Innovative Drugs Co., Ltd (Jiangsu, China). Racemic salbutamol sulfate (Rac-sal, >98%) and Dexamethasone Sodium Phosphate (Dex, 99%) were purchased from Shanghai Yuanye Bio-Technology Co., Ltd (Shanghai, China). Escherichia coli (0111: B4) LPS was purchased from Sigma-Aldrich (St. Louis, MO, USA). Ultra-pure water was produced by a Millipore A10 Milli-Q water purification system (Millipore, USA).

### Animals and treatments

Male BALB/c mice (6-8 weeks), weighing 20-25 g, were purchased from Guangdong Medical Laboratory Animal Center (Guangzhou, China) and housed in acrylic cages with free access to food and water under an environmentally controlled condition (room temperature: 25 ± 2 °C, humidity: 60 ± 5%, 12 h dark–light cycle). The experiments were approved by the Animal Ethics Committee of Guangdong University of Technology. All animal treatments were strictly in accordance with International Ethics Guidelines and the National Institutes of Health Guidelines Concerning the Care and Use of Laboratory Animals.

Mice were intraperitoneally injected with LPS (15mg/kg of body weight) to generate the LPS-induced mouse model of sepsis and lung injury. To examine the therapeutic effect of salbutamol enantiomers, mice were respectively treated with injection of R-sal low (0.1mg/kg), R-sal high (0.5mg/kg), S-sal (0.5mg/kg), or Rac-sal (1mg/kg) in saline at 30min, 6h, 12h post LPS challenge. For explore the protective effect of salbutamol enantiomers, mice were administered with R-sal low (0.1mg/kg), R-sal high (0.5mg/kg), S-sal (0.5mg/kg), or Rac-sal (1mg/kg) in saline twice per day for two days before LPS injection. Likewise, Dex (5mg/kg) was administered before or after LPS stimulus as comparison. The control group received equivalent normal saline instead.

### Survival rate

For survival rate, the mortality of each group of mice after LPS injection was monitored every 6h until 48h and then recorded every 24h until seven days.

### Sample collection and hematology analysis

Blood samples were collected six hours after LPS administration by intracardiac puncture under ether anesthesia. Whole blood samples were then partially transferred into EDTA-2K coated Eppendorf tubes. While serum was obtained after clotting for 2h at room temperature. Mice were then sacrificed and the lungs of mice were excised surgically and washed totally with ice-cold PBS to clear the blood and then blotted dry with filter paper. The left lung was fixed in 4% paraformaldehyde for histological examination. The right lung was stored at −80°C for MPO activity detection.

Whole blood samples were tested for the counts of total white blood cell (WBC) and differential WBCs (neutrophils, lymphocytes, monocytes and eosinophils) using a Procyte DX Hematology Analyzer (IDEXX Laboratories) in 4 h after blood collection.

### Lung histopathology

H&E staining assay was performed by Servicebio Inc. (Guangzhou, China). Briefly, left lung tissue samples were fixed in 4% paraformaldehyde overnight, followed by paraffin-embedded and then were sectioned at 4 μm for routine staining with hematoxylin and eosin. Images of the stained lung sections were obtained by an Axisplus image-capturing system (Zeiss, Germany) and then analyzed for lung inflammation and injury by pathologists blinded to grouping. Histological score was evaluated by a semi-quantitative scoring method as previously described[24]. Briefly, the overall lung injury score, including edema, inflammation, and hemorrhage, was graded on the following scale: normal (0), light (1), moderate (2), strong (3) and severe (4).

### Myeloperoxidase assay

Lung tissue was weighed and then homogenized in phosphate buffer (w/v: 1/5) with 1mM PMSF on crushed ice using a tissue grinder. The resulting mixture was then centrifuged at 12000×g for 10min at 4°C. The supernatant was then collected and measured using a mouse myeloperoxidase (MPO) ELISA kit following the manufacturer’s instructions (MultiSciences Biotech Co., Ltd, Zhejiang, China).

### Serum Lactate and cytokine assays

Serum samples were analyzed for the concentrations of Lactate using an enzyme-linked immunosorbent assay (ELISA) kit (Signalway Antibody, Maryland, USA) following the manufacturer’s protocol. The levels of serum tumor necrosis factor (TNF)-α, interferon (IFN)-γ, interleukin (IL)-6, IL-1β, IL-10 were assayed using a Luminex mouse pre-mixed Multi-Analyte kit according to standard instructions (LXSARM-06, R&D System, Minnesota, USA).

### Measurements of prognostic pulmonary function

Prognostic pulmonary function was assessed four days after LPS administration in conscious, freely moving, spontaneously breathing mice by a whole-body plethysmography (Date Sciences International, Inc., USA). After 2 min acclimation, mice were recorded for the values of respiratory frequency, tidal volume (TVb), minute volume (MVb) and enhanced pause (Penh) during a period of 3 min. In many cases, the degree of airway resistance was expressed as enhanced pause (Penh). The values of these parameters were automatically analyzed and calculated by the data acquisition software, FinePionte (Date Sciences International, Inc., USA).

### Statistical analysis

All data were presented as the means ± standard deviation (SD). The results were analyzed using GraphPad Prism software v8.4.2 (GraphPad, San Diego, CA, USA). Statistical comparisons were made by one-way ANOVA with post hoc Turkey multiple comparison tests. The data between two groups was compared by Student’s t test.

## Results

### Pre-treated with R-sal markedly improves the survival rate of LPS-induced septic mice and alleviates sepsis-induced acute lung injury

LPS is a widely used bacterial endotoxin to induce systemic inflammation and multiorgan dysfunction that resembles the initial clinical features of sepsis. First, we established a mouse model of experimental sepsis by intraperitoneally injection of a lethal dose of LPS (15mg/kg) and then monitored the mortality of each group of mice with salbutamol enantiomers, racemic mixture or Dex pretreatment for 7 days. As shown in Fig. 1A, LPS-treated septic mice showed a 48-h survival rate of 0%. In contrast, LPS mice pre-treated with R-sal high dose and Rac-sal, the mortality in these groups was completely inhibited. Meanwhile, LPS mice pre-treated with low dose of R-sal showed a survival rate of 60% at 48h, which was significantly increased when compared with LPS group. However, LPS mice pre-treated with S-ter and Dex showed no benefit to increase the survival rate of animals. Moreover, as compared with LPS-treated mice, mice pre-treated with Dex exhibited a significantly decreased survival rate (survival rate at 24h: 9.1% for Dex group *vs* 63.6% for LPS group; median survival time: 24h for Dex group *vs* 36h for LPS group), indicating that pretreatment with Dex have no protective effect and even accelerate death in LPS-induced septic mice. Histopathological analysis of lung sections by H&E staining showed that the lungs of LPS-treated mice exhibited features of damage and acute inflammation, including inflammatory cell infiltration, thickened alveolar septa and congestion. With pretreatment of R-sal and double molar Rac-sal, the degree of these damages was ameliorated respectively. However, semiquantitative overall histological scores revealed that only high dose of R-sal significantly attenuated lung damages compared to LPS-treated group. S-sal or Dex pretreatment to some extent had a risk of augmented alveolar exudates and congestion/hemorrhage but with no significant difference when compared with the LPS group.

**Fig. 1.**
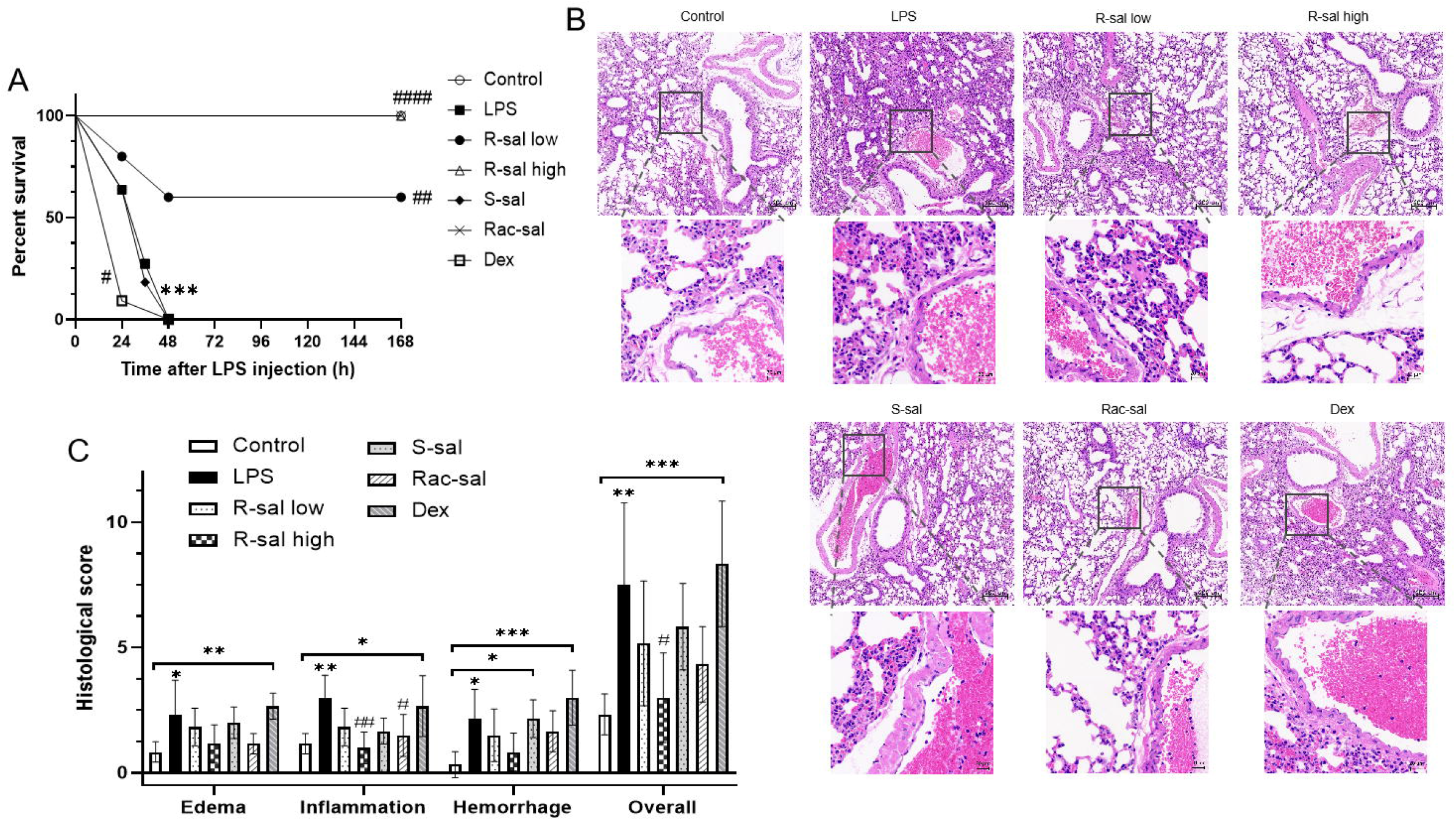
Effects of R-sal, S-sal, Rac-sal or Dex pretreatment on the survival rate and lung damages in LPS-induced septic mice. Each ten mice were pre-treated with R-sal low dose, R-sal high dose, S-sal, or Rac-sal, or saline twice per day for two days and then intraperitoneally injected with either LPS (15mg/kg) or isometric saline. (A) 7-day survival curves of each group of mice. Survival curves were calculated according to the Kaplan-Meier method. Survival analysis was performed using Log-Rank test; (B) Hematoxylin and eosin staining of lung sections was assessed at 6h after LPS injection; (C) Histological scores. Statistical data is shown as mean ± SD (n = 6). *p<0.05, **p<0.01, ***p<0.001 vs the control group; ^#^p<0.05, ^##^p<0.01, ^###^p<0.001, ^####^p<0.0001 vs the LPS group.

### Pre-treated with R-sal reduces MPO level in lung tissue of LPS-induced septic mice

To further investigate the protective effect of R-sal, MPO determination was conducted to assess the degree of lung lesions. As shown in Fig.2, the MPO expression markedly increased in LPS-treated mice, which could suppress by R-sal in a dose-dependent manner. Administration with R-sal high dose significantly inhibited the increased MPO level in the lung of septic mice.

**Fig. 2.**
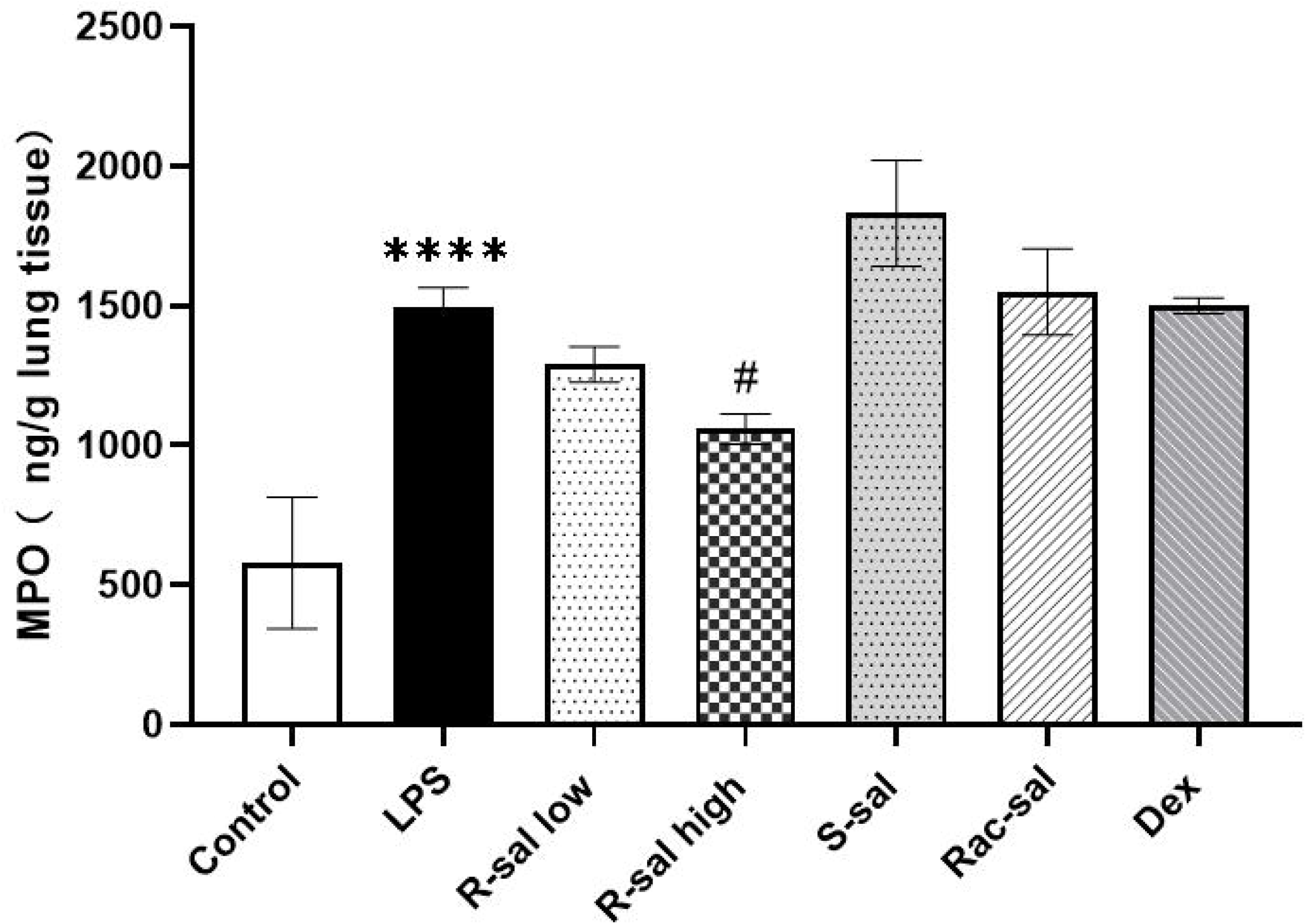
Effects of R-sal, S-sal, Rac-sal or Dex pretreatment on myeloperoxidase (MPO) level in the lung tissues of LPS-induced septic mice. The lung tissue for the MPO determination was collected 6h after LPS challenge. Data are presented as mean ± SD (n=3). ****p<0.0001 vs the control group; ^#^p<0.05 vs the LPS group.

### R-sal pretreatment restores lymphocytes, significantly suppresses the percentage of monocytes and serum lactate

Depleted lymphocytes and elevated lactate level are associated with pathophysiology of sepsis and sepsis-induced complications[1][25]. As shown in Fig. 3A, at 6h after LPS injection, total WBCs in LPS-induced septic mice was drastically reduced when compared with the normal saline treated mice. It has found that the significantly reduced white blood cell is lymphocyte according to the differential WBCs counts (Fig. 3B). Pretreatment with R-sal dose dependently restored the decreased lymphocytes in mice after LPS exposure. Meanwhile, R-sal or Rac-sal pretreatment, not S-sal, could substantially mitigated the elevated MONO% in LPS-induced septic mice (Fig. 3C). As show in Fig. 3D, serum lactate concentration was markedly increased in mice treated with LPS when compared with the control group, which could significantly inhibit by R-sal high dose, double molar amount of Rac-sal and Dex.

**Fig. 3.**
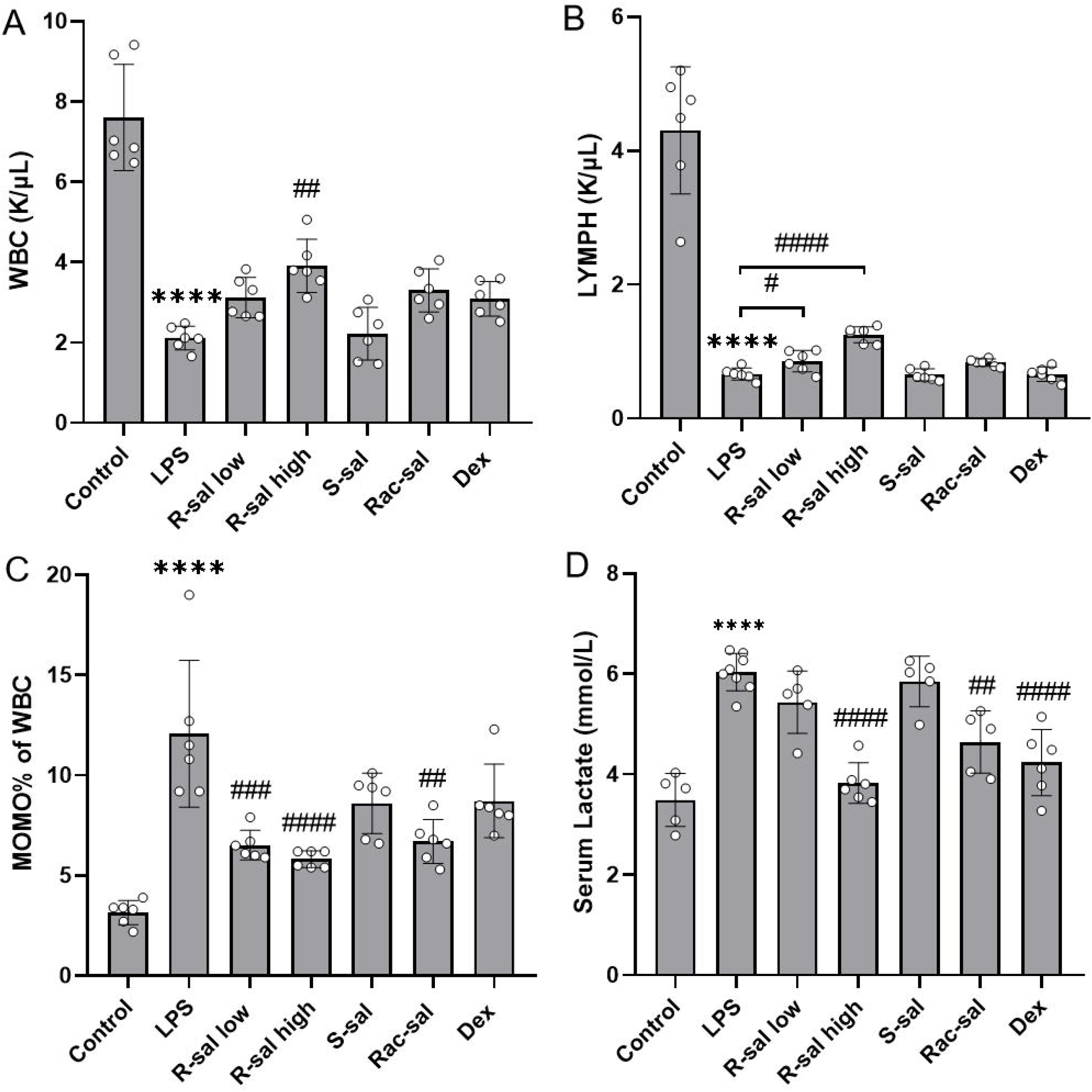
Effects of R-sal, S-sal, Rac-sal or Dex pretreatment on total white blood cells (WBCs), differential WBCs and serum lactate in LPS-induced septic mice. (A) WBC counts; (B) Lymphocytes counts; (C) The percentage of monocytes in blood; (D) The level of serum lactate. Data are presented as mean ± SD (n=3). ****p<0.0001 vs the control group; ^#^p<0.05, ^##^p<0.01, ^###^p<0.001, ^####^p<0.0001 vs the LPS group.

### R-sal pretreatment significantly suppresses systemic inflammatory cytokines in LPS-induced septic mice

A progressive secretion of inflammatory cytokines, like TNF, IL-1β and IL-6, is involved in the development of sepsis and sepsis-induced multiorgan dysfunction. As shown in Fig.6, after 6h of LPS injection, mice displayed a robust increased serum concentration of both pro-inflammatory cytokines IFN-γ, TNF-α, IL-1β, IL-6 and anti-inflammatory cytokine IL-10. Pretreatment with R-sal and double molar of Rac-sal reduced LPS-induced production of IL-1β and markedly decreased LPS-induced production of IFN-γ, TNF-α and IL-6. Whereas, S-sal treatment was found no reduction in LPS-induced production of inflammatory cytokines. Notably, mice pre-treated with S-sal could activate the release of IFN-γ, TNF-α and IL-1β in system, implicating a bad role of S-sal in host immunoregulation. Dex pre-treated mice exhibited less concentrations of IFN-γ and IL-1β but not IL-6 than those in LPS-treated mice and a significantly augmented TNF-α level compared to LPS group. In addition, high dose of R-sal pretreatment significantly increased the concentration of anti-inflammatory cytokine IL-10 when compared with the LPS group.

### R-sal pretreatment restores the prognostic lung function of survived septic mice

To further investigate the therapeutic effects of R-sal and Rac-sal in protecting sepsis-induced lung injury, we assessed the prognostic lung function of survived septic mice by means of noninvasive in vivo plethysmography. As shown in Fig. 5, the Penh and minute volume (Mvb) of mice treated with R-sal high dose showed no significant difference relative to those of mice in the control group, indicating a complete recovery of respiratory function of septic mice by R-sal high dose pretreatment. Double molar amount of Rac-sal pretreatment showed an increased tendency of increased Penh without significant difference and significantly decreased Mvb in comparison with the control group.

**Fig. 4.**
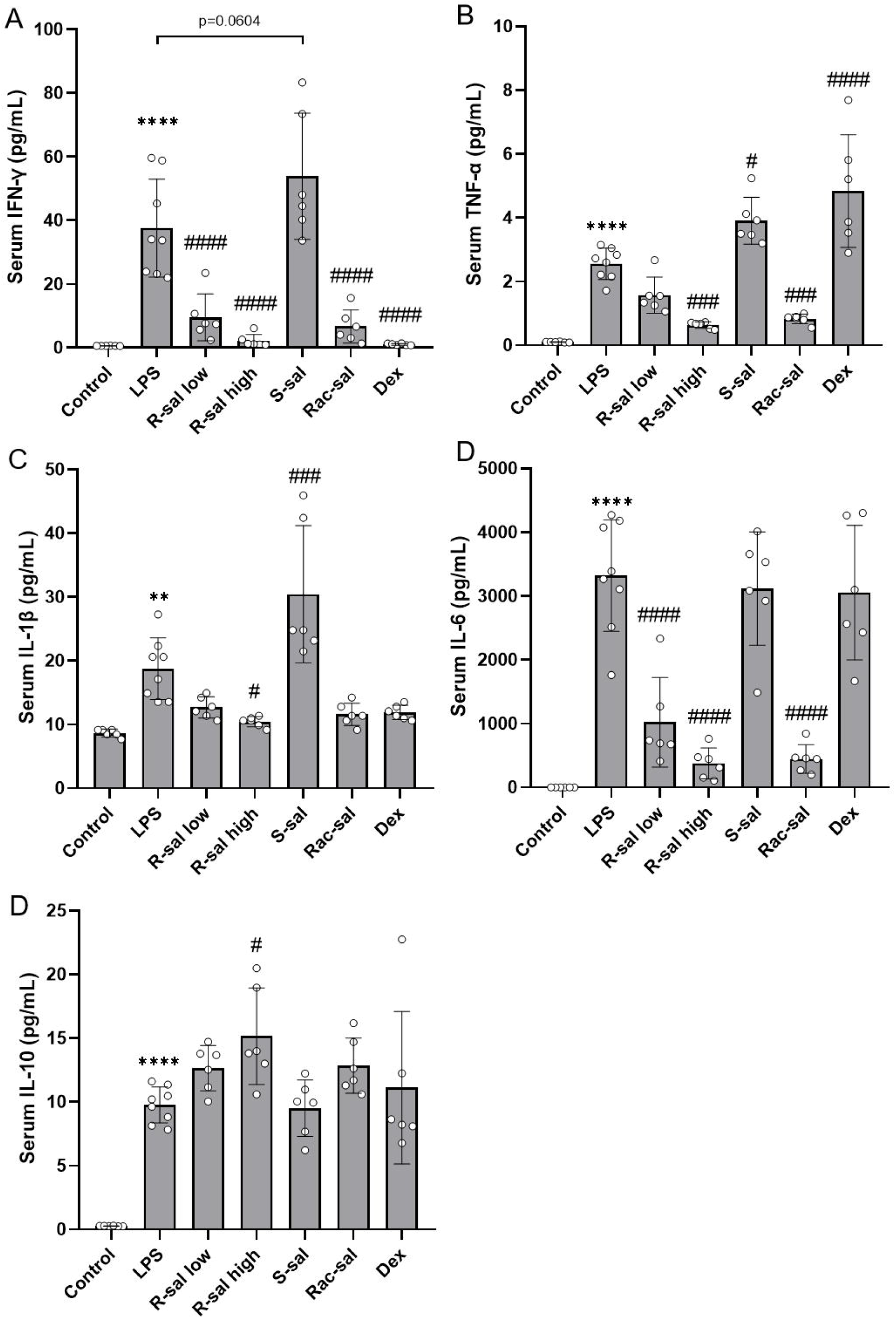
Effects of R-sal, S-sal, Rac-sal or Dex pretreatment on systemic inflammatory cytokines in LPS-induced septic mice. (A) serum IFN-γ; (B) serum TNF-α; (C) serum IL-1β; (D) serum IL-6; (E) serum IL-10. Data are presented as mean±SD (n=6-8). **p<0.01, ****p<0.0001 vs the control group; ^#^p<0.05, ^###^p<0.001, ^####^p<0.0001 vs the LPS group.

**Fig. 5.**
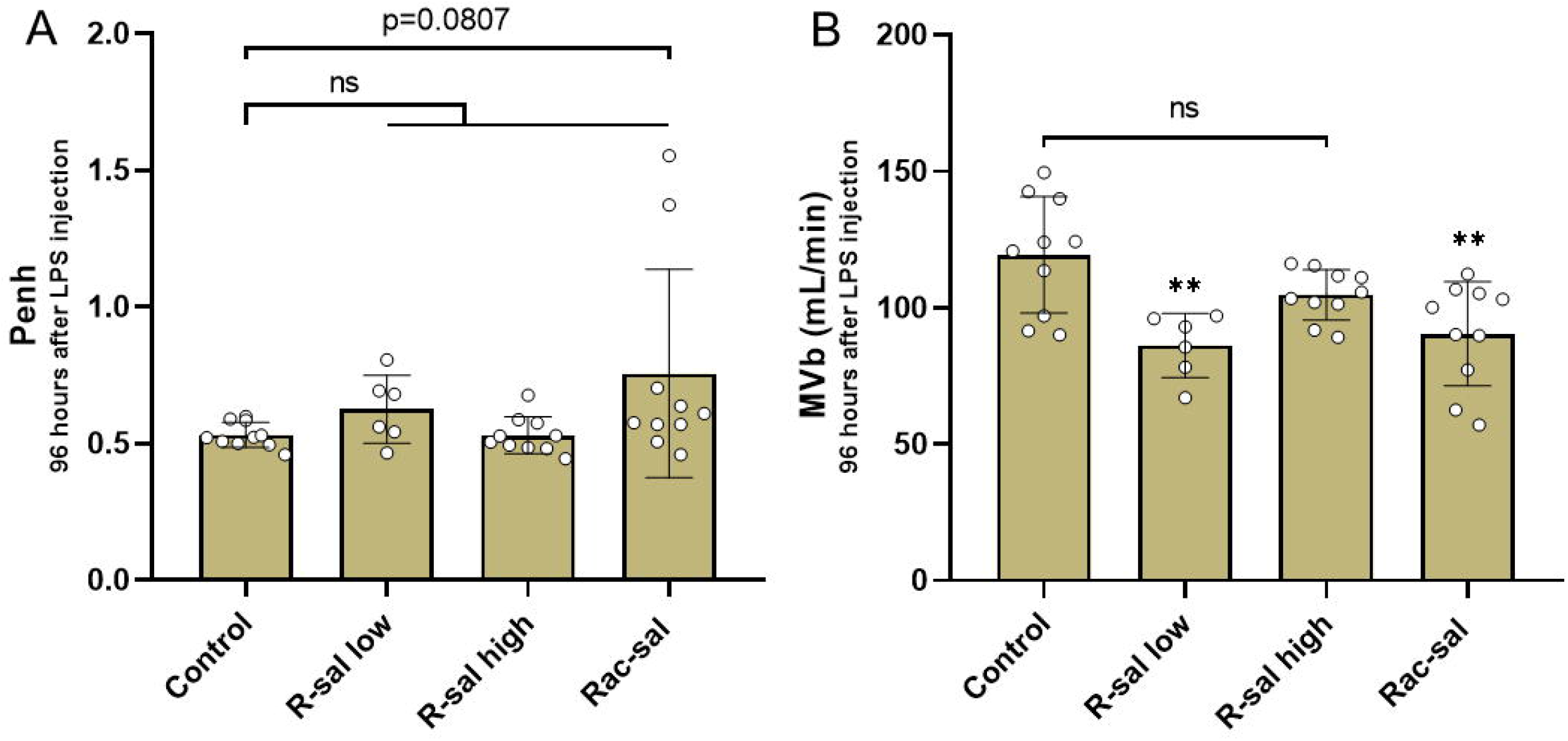
Effects of R-sal and Rac-sal pretreatment on the prognostic lung function of survived septic mice. Each ten mice were pre-treated with R-sal low dose, R-sal high dose, S-sal, or Rac-sal, or saline twice per day for two days and then intraperitoneally injected with either LPS (15mg/kg) or isometric saline. The lung function of the survived septic mice in each group was assessed by means of noninvasive in vivo plethysmography on day 4. (A) The values of Penh; (B) Minute volume; Data are presented as mean ± SD (n=6-8). **p<0.01 vs the control group.

### Treated with R-sal (0.5mg/kg) also markedly improves the survival rate of LPS-induced septic mice

Considering the clinical practice, we also examine the effect of R-sal on septic mortality when administrated at 30min, 6h and 24h after LPS stimulation. As presented in Fig. 6, Kaplan-Meier survival curves revealed that mice with LPS-induced sepsis showed a 7-day survival rate of 30%. R-sal treatment dose-dependently inhibited the mortality of septic mice. R-sal high dose significantly increased the 7-day survival rate to 90% while Rac-sal exhibited a declined survival rate to 80%. As Noted, Dex administration after LPS challenge demonstrated to markedly improves the survival rate of septic mice to 100% (7-day), which was inconsistent with the result of Dex pretreatment.

**Fig. 6.**
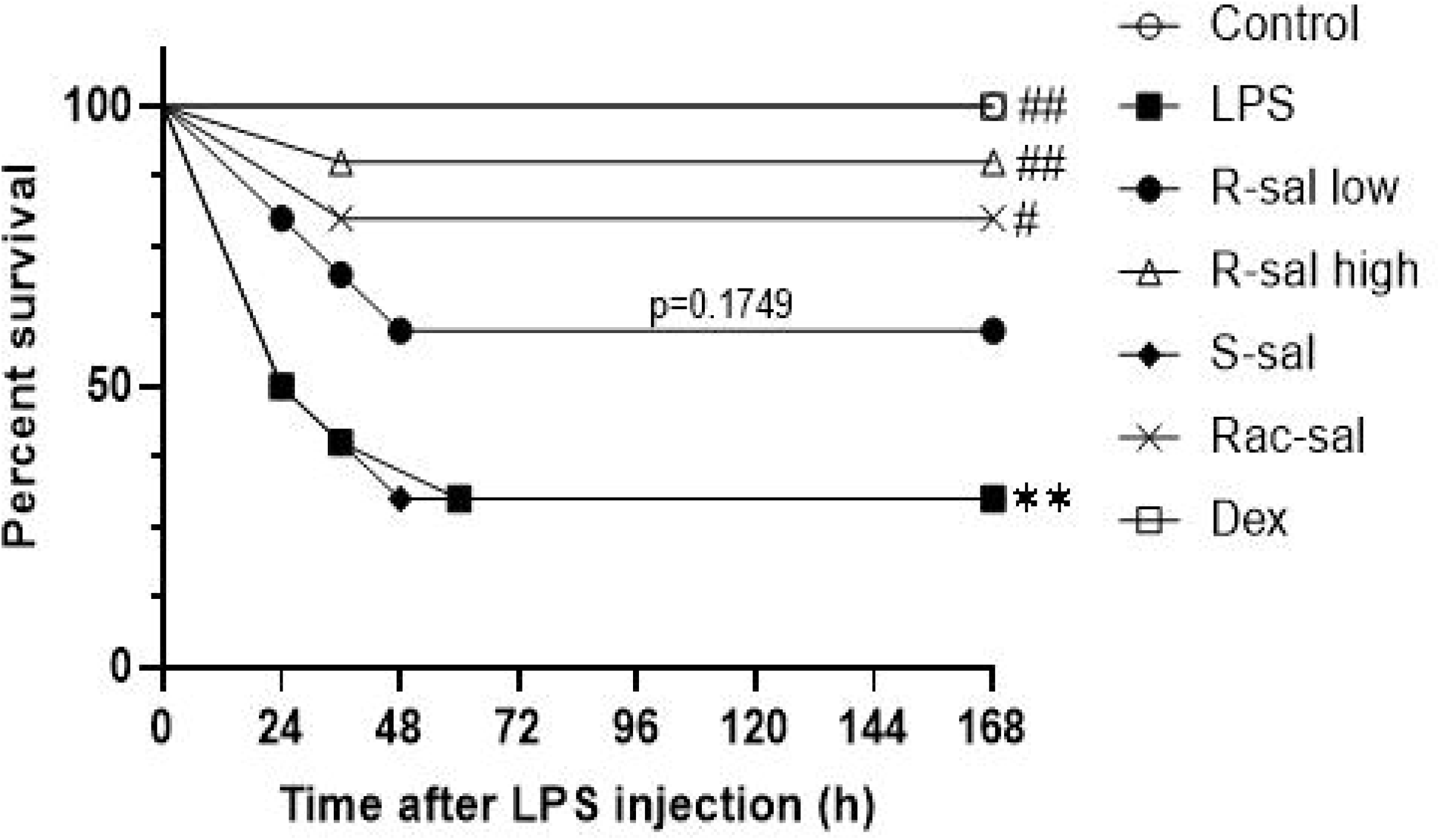
Seven-day survival rate of LPS-induced septic mice treated with R-sal, S-sal, Rac-sal or Dex. Each ten mice were intraperitoneally administered with either LPS (15mg/kg) or isometric saline, followed by R-sal, S-sal, Rac-sal or Dex, or saline treatment at 30min, 6h, and 24h. Survival curves of septic mice were calculated according to the Kaplan-Meier method. Survival analysis was performed using log-rank test. **p<0.01 vs the control group; ^#^p<0.05, ^##^p<0.01 vs the LPS group.

### Effects of R-sal (0.5mg/kg) administration on normal mice

To identify whether R-sal (0.5mg/kg) had any harmful effects on normal mice, we conducted several representative examinations. As shown in Fig. 7, treatment of mice with high dose of R-sal for three days have no significant effect on the lung architecture, lung function, the level of serum lactate and serum inflammatory cytokines.

**Fig. 7.**
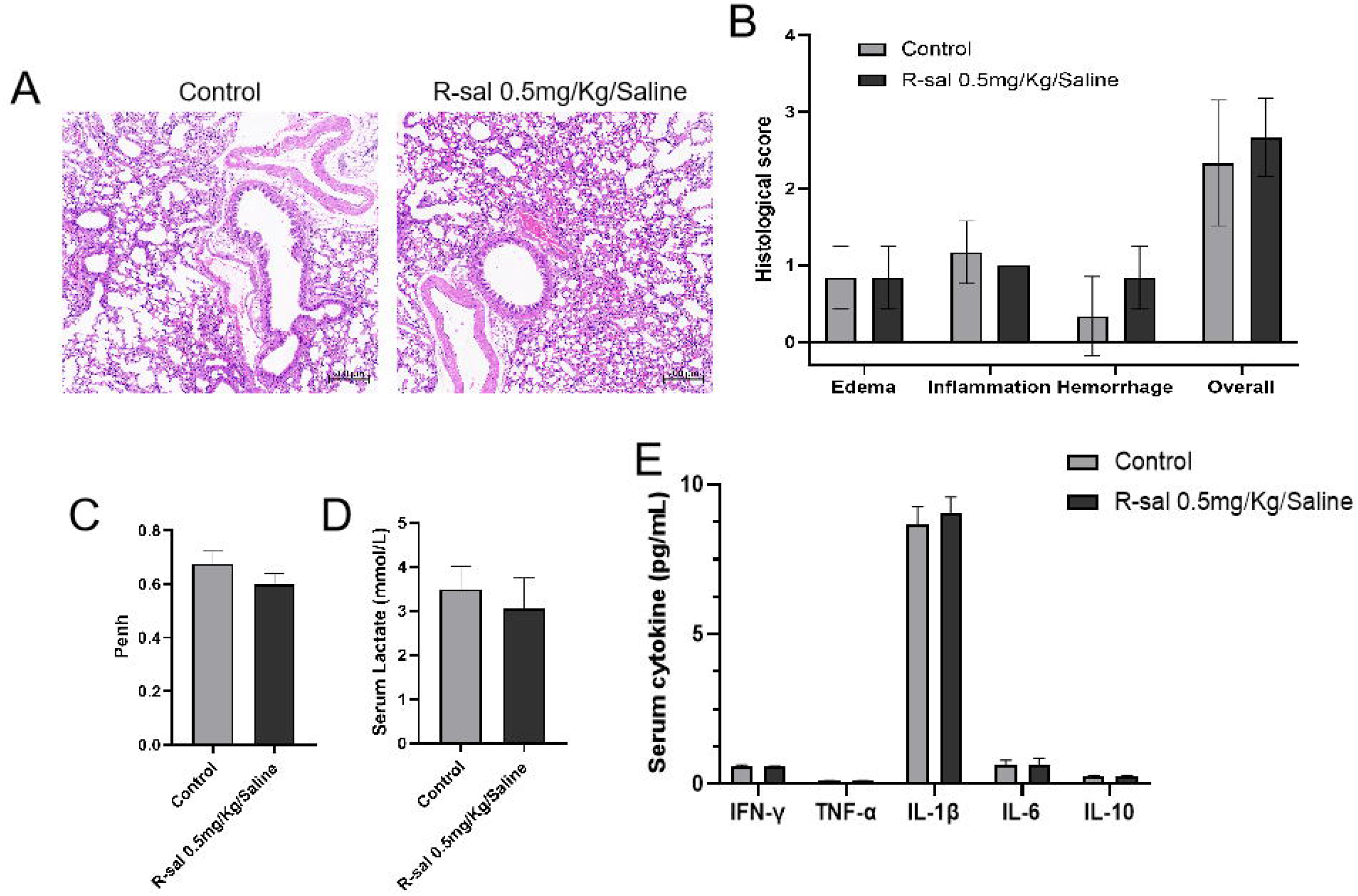
Effects of R-sal (0.5mg/kg) administration on normal mice. Mice were treated with intraperitoneal injection of R-sal (0.5mg/kg) twice per day for three consecutive days. On day 3, the lung function (C) of each group of mice were assessed by means of noninvasive in vivo plethysmography. Then the lungs and serum samples were collected under anesthesia. (A) hematoxylin and eosin staining; (B) Histological scores; (D) the level of serum lactate; (E) the levels of serum cytokines. Data are presented as mean±SD (n=6-10).

## Discussion

Sepsis is a life-threatening disease with relatively high mortality. In clinical practice, elimination of the infection and haemodynamic stabilization are the mainstay for the prevention and management of sepsis. In addition, immunomodulatory therapy has been gained increasingly attention for the treatment of sepsis, although limited effective medications have been found. Salbutamol, which consists of R-enantiomer and S-enantiomer, is a broadly used short-acting β2-agonist for asthma and obstructive lung diseases. A growing body of evidences have revealed the effects of β2-agonists on possessing anti-inflammatory properties on airways and reducing pro-inflammatory mediators as well as preventing tissue oedema and exudate. Despite sufficient researches have demonstrated the compelling protective effects of salbutamol and other β2-agonists on preventing sepsis in pre-clinical experiments, these agents have not been shown to improve outcomes in clinical trials in ARDS (a syndrome in critically ill sepsis)[26][27]. As we acknowledged, two enantiomers of salbutamol exhibit differential pharmacological effects. Accumulating evidences and our previous studies have proven the superiority anti-inflammatory effect of R-salbutamol and the pro-inflammatory effect of S-enantiomer[28][29]. Therefore, in the current study, we utilize a mouse model of experimental sepsis to investigate the differential effects of R-sal, S-sal and Rac-sal on the survival rate, lung and systemic inflammation in mice. Our results demonstrated the superiority of R-sal in regulating the LPS-induced immune dysfunction of sepsis in mice.

LPS has been widely used to induce systemic inflammation and multiorgan dysfunction in mice to mimic the clinical features of sepsis. Lethal septic shock is required to examine survival following sepsis. We first established a mouse model of experimental sepsis by intraperitoneally injection of a lethal dose of LPS (15mg/kg). Mice with LPS challenge result in a 48h-survival rate of 0%. R-sal pretreatment, as well as Rac-sal, dramatically improved the survival rate to 100% while mice pre-treated with S-sal exhibited a slight accelerated lethality with no significant difference in comparison with the LPS group. Unexpectedly, we found that, in this setting of sepsis, Dex pretreatment showed an deteriorated lethality of mice (Fig.1A). However, administration of Dex after LPS injection presented a marked increased survival rate of 100% (Fig.6), which in consistent with the previous reports[30]. These results indicate a toxic effect of Dex pretreatment might due to the suppression of immune system. As presented in Fig.6, R-sal high dose significantly increased the 7-day survival rate to 90%, suggesting the potential therapeutic effect of R-sal. Whereas, double molar amount Rac-sal exhibited a diminished survival rate of 80%, possibly ascribed to the S-enantiomer.

Sepsis-induced lung injury is one of the severest complications in sepsis. Lung damage in sepsis is caused by excess or severe inflammation. β2-agonists have been reported to ameliorate both the endotoxemia and cecal ligation and puncture (CLP)-induced sepsis-associated organ injury through regulating inflammatory cytokines release[20][26]. In similar with the earlier researches, our results demonstrated that R-sal exerted a more effective attenuation of lung inflammation, edema and congestion in LPS-induced septic mice than double molar Rac-sal. In contrast, Dex pretreatment showed a tendency of increased alveolar exudates and hemorrhage in the lungs of septic mice which might lead to the accelerated lethality in the Dex group (Fig.1B). In the setting of sepsis, lung MPO activity can reflect the degree of tissue infiltration by neutrophils, which directly contributes to the injury of the lungs. In this regard, our findings showed that pre-treated with R-sal high dose significantly suppressed the augmented MPO level in the lung of LPS-induced septic mice (Fig.2).

Sepsis is characterized not only by increased inflammation but also by immune suppression[31]. Depleted lymphocytes and elevated lactate level are associated with pathophysiology of sepsis and sepsis-induced complications[25]. In the present study, whole white blood cells, mostly lymphocytes were drastically reduced and the percentage of monocytes was significantly increased six hours after LPS stimulation. R-sal pretreatment dose-dependently impeded the reduction of lymphocytes and the elevated percentage of monocytes. In contrast with R-sal high dose group, double molar Rac-sal exhibited a waning efficacy of protection (Fig.3A, 3B and 3C). Literatures report that LPS can be detected via. Pathogen-associated molecular patterns (PAMPs) by monocytes and macrophages to induce cytokine production and other important responses[32]. In turn, we found that R-sal high dose pretreatment effectively suppressed the pro-inflammatory cytokines IFN-γ, TNF-α, IL-1β and IL-6 and increased the anti-inflammatory cytokine IL-10. Conversely, S-sal showed an aggravated release of IFN-γ, TNF-α and IL-1β in septic mice (Fig.4). In a report by Bosmann et. al.[21], S-sal, but not R-sal, showed an anti-inflammatory effect in ALI mouse model, which is inconsistent with our findings. However, accumulating evidences have demonstrated R-enantiomer of salbutamol exerts the anti-inflammatory effect whereas S-sal to some extent can generate pro-inflammatory effect. In spite of the potent effectiveness of R-sal, β2-agonists is thought to activate β2-adrenorecepotor of skeletal muscle and then give rise to lactic acidosis, which limits the clinical application of β2-agonists for sepsis [33][34]. However, in a case report suggests that the lactic acidemia caused by β2-agonists is not associated with hypoxia or shock[35]. Nonetheless, in our experimental model of sepsis, pretreatment of both R-sal and Rac-sal could inhibit the serum lactic acidosis of septic mice (Fig.3D).

Patients who survived form sepsis are more prone to suffer an long-term organ damage and disability, resulting in a poor prognostic of disease. Pulmonary function could be a effective measure to assess the recovery of diseases[1][36].In our results, the minute volume of septic mice pre-treated with R-sal high dose returned to normal, together with a lower value of Penh. In contrast, Rac-sal exhibited a less potential effect on the recovery of lung function (Fig.5). Furthermore, we found that normal mice administrated with R-sal high dose for three days indicated no significant changes in lung architecture and function, hematology and inflammatory mediators (Fig.7). Collectively, our findings demonstrated that pre-treated or treated with R-sal high dose could markedly improve the survival rate of septic mice and confirmed that R-sal exhibited a better efficacy in regulating immune dysfunction of sepsis than its racemic mixture.

In spite of the effective immune regulation effect of R-sal, β2-agonist are always associated with a typical side effect, dose-related tachycardia, which should be take into consideration[37]. However, the randomized controlled trials in pediatric clearly report that there is an advantage of using R-sal over Rac-sal in reducing tachyarrhythmias[38]. Moreover, R-sal at a lower dose could have a potential in reducing its cardiotoxicity side effects. Additionally, an oral dose of 25mg/d salbutamol in pregnant woman for prevention of preterm labor has been reported by Motazedian[39]. Most importantly, selective stimulation of β2-adrenoreceptor is reported to antiapoptotic rather than cardiotoxicity. And enough evidence has been accumulated in the literature to propose particularly β2 activation and β1 blockade as new promising therapeutic targets for sepsis[40][41]. But it still needs to further investigate as to whether R-sal could show a beneficial effect over racemate on sepsis in clinical practice.

## Conclusion

In conclusion, the present study found that pre-treated or treated with R-sal high dose could markedly improve the survival rate in an LPS-induced lethal mouse model of sepsis. Dex pretreatment showed a deteriorated mortality of mice, which might alert to the potential risk of Dex administration before infection. R-sal pretreatment exhibited a better efficacy than rac-sal in reduction of sepsis-induced lung damage in mice, accompanying by lower MPO level, serum lactate and recovered prognostic lung function. R-sal pretreatment significantly restored the lymphocytes and suppressed the percentage of monocytes. We demonstrated R-sal effectively inhibited the pro-inflammatory cytokines release and increased anti-inflammatory IL-10 level. Whereas S-sal could aggravate the release of IFN-γ, TNF-α and IL-1β. Collectively, our findings suggest that R-sal is a promising immune regulator for sepsis and sepsis-related lung injury in clinic.

## Acknowledgement

This work was supported by the National Science and Technology Major Projects for “Major New Drugs Innovation and Development”, 2019ZX09301120.

